# Empagliflozin targets a renal neuro-epithelial-immune axis in heart failure

**DOI:** 10.64898/2026.03.31.715595

**Authors:** Jennifer N. Coelho, Livia C. Simonete, Joao Carlos Ribeiro-Silva, Érika F. Jesus, Andreia Boaro, Flavia L. Martins, Jose Wilson Correa do Nascimento, Larissa Ferreira dos Santos, Danubia Silva dos Santos, Ednei L. Antonio, Andrey J. Serra, Adriana C. C. Girardi

**Affiliations:** Departamento de Cardiopneumologia, Faculdade de Medicina, Universidade de São Paulo, São Paulo, SP, Brazil; SUNY Upstate Medical University, Syracuse, NY, USA; Department of Physiological Sciences, Biological Sciences Institute, Federal University of Amazonas, Manaus, Brazil; Division, Department of Medicine, Federal University of São Paulo, São Paulo, São Paulo, Brazil

**Keywords:** SGLT2 inhibitors, heart failure, norepinephrine, inflammation, kidney, proximal tubule

## Abstract

**Background:** Persistent neurohormonal activation is a key driver of maladaptive remodeling and disease progression in heart failure (HF). Sodium-glucose cotransporter 2 inhibitors (SGLT2is) confer robust renoprotective effects in HF; however, the extent to which these benefits involve modulation of renal neurohormonal activity remains unclear. We hypothesized that SGLT2i-mediated renoprotection in HF is associated with attenuation of excessive renal neurohormonal activation.

**Methods:** Male rats with myocardial infarction-induced HF and sham controls were fed standard chow or chow containing empagliflozin (EMPA, 300 mg/kg) for four weeks, followed by assessment of renal inflammatory and neurohormonal markers. Parallel in vitro studies in THP-1 macrophages and HK-2 proximal tubule cells evaluated the direct effects of EMPA on norepinephrine (NE)-dependent tubular inflammatory signaling.

**Results:** HF rats displayed higher renal cortical renin gene expression and angiotensin II concentrations, which remained unaffected by EMPA. Conversely, EMPA normalized the elevated urinary NE excretion and renal cortical NE content observed in HF rats. Given the inflammatory role of sympathetic hyperactivity, we assessed renal macrophage polarization. EMPA-treated HF rats showed reduced expression of pro-inflammatory markers (*Tnf, Ccr2, Nos2, Il-6*) and increased expression of markers associated with a reparative macrophage profile (*Arg1, Mrc1, CD163*), supported by higher CD206⁺ macrophages in kidney sections. While EMPA did not directly alter THP-1 macrophage activation *in vitro*, it significantly reduced NE-induced SGLT2 expression and interleukin-6 (IL-6) release by HK-2 human proximal tubule epithelial cells.

**Conclusion:** These findings support a model in which SGLT2 inhibitors confer renoprotection in HF by suppressing renal sympathetic hyperactivity, independently of the intrarenal renin-angiotensin system, thereby disrupting a maladaptive renal neuro-epithelial-immune axis and promoting a reparative macrophage phenotype.

**CLINICAL PERSPECTIVE:** *What’s new?:*

- This study identifies a renal neuro-epithelial-immune axis underlying empagliflozin-mediated renoprotection in heart failure.
- Empagliflozin reduces renal cortical and urinary norepinephrine levels in heart failure without altering intrarenal renin-angiotensin system activity, revealing a distinct neurohumoral target of SGLT2 inhibition.
- This sympatholytic effect is associated with a shift in renal macrophages toward a reparative (M2) phenotype, without changes in total macrophage abundance.
- Empagliflozin blocks norepinephrine-induced SGLT2 upregulation, limiting proximal tubular glucose reabsorption and IL-6 production, and linking sympathetic signaling to renal inflammation.

*What are the clinical implications?:*

- Our findings provide a mechanistic basis for the additive cardiorenal benefits of SGLT2 inhibitors in heart failure, beyond conventional RAS-directed therapies.
- Targeting renal sympathetic-driven inflammation may help preserve kidney function and attenuate the progression of cardiorenal syndrome.
- Suppression of a renal neuroinflammatory pathway may help explain the early and sustained benefits of SGLT2 inhibitors across heart failure phenotypes, including nondiabetic patients.

## INTRODUCTION

Heart failure (HF) is a complex clinical syndrome caused by structural and/or functional cardiac abnormalities that impair ventricular filling and/or ejection^1^, often leading to secondary renal dysfunction that culminates in the development of cardiorenal syndrome (CRS)^2^.

This inter-organ crosstalk is driven by a complex neurohumoral milieu: activation of the renin-angiotensin-aldosterone system (RAAS), heightened sympathetic nervous system (SNS), and increased arginine vasopressin (AVP) signaling. Together, these pathways promote sodium and water retention, enhance renal vasoconstriction, and reduce renal perfusion, ultimately worsening kidney dysfunction and perpetuating congestion. While counter-regulatory mechanisms (most notably natriuretic peptides) attempt to restore volume homeostasis, they are insufficient in advanced HF^3–5^. Simultaneously, renal sensory (afferent) signaling and impaired baroreflex control may amplify central sympathetic outflow, creating a maladaptive feed-forward mechanism linking systemic neurohumoral activation to intrarenal hemodynamic and tubular dysfunction^6–8^.

Originally developed as glucose-lowering agents, the sodium-glucose cotransporter 2 inhibitors (SGLT2is) have become a cornerstone of HF therapy^9,10^. Clinical evidence demonstrates that SGLT2is significantly reduce cardiovascular mortality and HF hospitalizations across the spectrum of ejection fraction^11–14^ beyond their glycemic effects. These benefits derive, at least in part, from the cardioprotective and renoprotective effects of SGLT2is. The cardioprotective effects of SGLT2is are associated with modulation of intracellular calcium and sodium levels, leading to reduced arrhythmias and attenuation of right ventricular remodeling^15–18^. On the other hand, the renoprotective effects are associated with the promotion of natriuresis, slowing the decline in the glomerular filtration rate (GFR), reduction of albuminuria, and attenuation of renal structural remodeling^19–24^. However, the precise molecular mechanisms underlying the benefits of SGLT2is in HF remain incompletely understood.

One emerging mechanism is the potential for SGLT2is to attenuate peripheral SNS activity in HF. Chronic sympathetic hyperactivity drives adverse cardiac remodeling, pathological hypertrophy, and impaired contractility, effects that are mitigated by attenuation of sympathetic activity. In the kidneys, HF is associated with attenuated sympathoinhibitory responses to acute volume expansion, compromising fluid homeostasis^25,26^.

We have previously shown that the SGLT2 inhibitor empagliflozin (EMPA) restores diuretic and natriuretic responses to acute volume expansion in rats with heart failure and attenuates SGLT2 and NHE3 upregulation in the renal proximal tubule. These effects parallel the functional benefits of renal denervation, supporting a role for modulation of renal sympathetic tone in EMPA-mediated renoprotection^24,26^. Building on these findings, the present study sought to determine whether SGLT2 inhibition affects renal neurohormonal activity in HF. We tested the hypothesis that the renoprotection by SGLT2is in HF is associated with the attenuation of excessive renal neurohormonal activation and the subsequent reduction of renal inflammation.

## METHODS

### Rats

All animal experiments were conducted in compliance with the National Council for the Control of Animal Experimentation (CONCEA) guidelines and approved by the Ethics Committee on Animal Use of the University of São Paulo Medical School (Protocol #941/2018). Male Wistar rats (7 weeks old, 190–220 g) were obtained from the Institute of Biomedical Sciences, University of São Paulo, and randomly assigned to sham surgery or myocardial infarction (MI) induced by ligation of the left anterior descending (LAD) artery, as previously described^27^. Rats were anesthetized with intraperitoneal injection of ketamine (50 mg/kg) and xylazine (10 mg/kg) and mechanically ventilated. A thoracotomy was performed, and the LAD was ligated ∼3 mm from its origin using 6-0 Prolene suture. Sham-operated rats underwent the same procedure except for LAD ligation. Following surgery, rats were housed under controlled conditions with food and water *ad libitum*. Four weeks after surgery (randomization), HF was confirmed in surviving MI rats based on serum brain natriuretic peptide (BNP) levels (> 1.0 ng/ml) and fractional area change (FAC, <40%) determined by echocardiography^24^. Echocardiography was performed under 1.5% isoflurane anesthesia using the Sonos 5500 ultrasound equipment with a 12–1 (4MHz transducer)^24^. FAC was calculated as [LV end-diastolic area (LVEDA) - LV end-systolic area (LVESA) / LVEDA] × 100%. Serum BNP levels were measured by enzyme-linked immunosorbent assay (ELISA; BNP 32 Rat ELISA kit, ab108815, Abcam) according to the manufacturer’s instructions. Echocardiography and BNP assessments were conducted blinded to group assignments. HF (n=32) and sham (n=32) rats were randomized to receive EMPA (300 mg/kg of chow) (A12440, Adooq Bioscience, Irvine, CA) or no treatment for four weeks. At the study endpoint (posttreatment), rats were anesthetized with isoflurane (5% in O_2_-enriched air; maintenance dose 3%) and the lungs, heart, kidneys, and arterial blood (∼1.5 ml from the abdominal aorta) were collected. The study design is depicted in **Figure 1A**.

**Figure 1.**
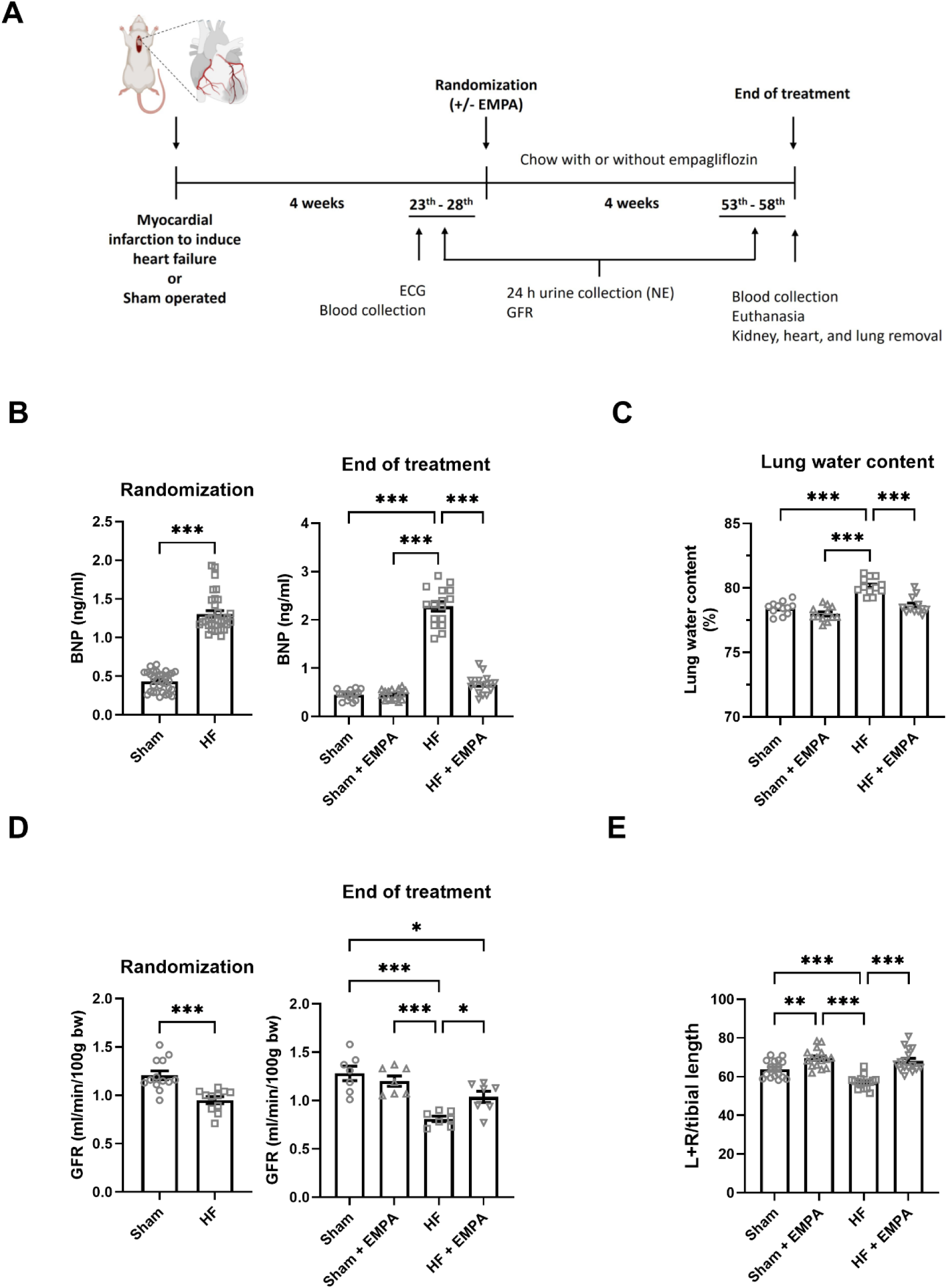
Empagliflozin attenuates congestion and preserves renal function in rats with heart failure. **(A)** Study timeline. Male Wistar rats underwent sham surgery or myocardial infarction (MI) by left anterior descending coronary artery ligation to induce heart failure (HF). Four weeks after surgery (randomization), animals received empagliflozin (EMPA, 300 mg/kg chow) or not for an additional 4 weeks. Glomerular filtration rate (GFR) was measured transcutaneously using FITC-sinistrin at randomization and posttreatment. **(B)** Plasma brain natriuretic peptide (BNP) concentrations were measured at randomization and at the end of the treatment period. **(C)** Lung water content. **(D)** Glomerular filtration rate (GFR) at randomization and at the end of the treatment period. **(E)** Kidney weight-to-tibia length ratio. Data are presented as mean ± SEM. Statistical analyses were performed using an unpaired Student’s *t* test for randomization data (panels B and D) and two-way ANOVA followed by Tukey’s post hoc test for posttreatment comparisons (panels B–C and E; *n* = 16 rats per group; panel D, *n* = 7 rats per group). *P < 0.05, **P < 0.01 and ***P < 0.001.

### Biometric and morphometric analysis

Lungs and kidneys were immediately excised and weighed. Kidney weights were normalized to left tibia length. Lung samples were dried at 70°C for 48h, and lung water content was determined using the formula: % Lung Water Content = [(Wet Weight − Dry Weight) / Wet Weight] × 100. The kidneys were promptly harvested for protein isolation, immunohistochemistry, or RNA extraction.

### Transcutaneous measurement of glomerular filtration rate (GFR)

GFR was assessed on days 25–26 (randomization) and 55–56 (posttreatment) (**Figure 1A**) using the MediBeacon GmbH system (Mannheim, Germany) and following a modified method described by Schreiber et al.^28^ Briefly, rats were anesthetized with isoflurane (5% in oxygen-enriched air for induction; 3% for maintenance). The fur over the right flank (between the rib cage and iliac crest) was shaved using an electric trimmer, and a sensor was secured to the exposed skin with adhesive tape to ensure stability. A 5-minute stabilization period was observed while the rats’ tails were submerged in a 40°C water bath to dilate the caudal vein. Fluorescein isothiocyanate (FITC)-sinistrin (7.5 mg/100 g body weight) (FTC-FS001, Fresenius Kabi GmbH, Austria) was injected into the caudal vein. Rats were then placed in individual cages without food or water for a 2-hour recording period, after which the sensor was removed, and the rats were returned to normal housing conditions. GFR data were analyzed using the MB Studio 3 Software (MediBeacon).

### Determination of renal Angiotensin II, Angiotensin (1-7), Interleukin-6, and urinary norepinephrine

Renal cortical concentrations of Angiotensin II (Ang II), Angiotensin-(1–7) [Ang-(1–7)], and Interleukin-6 (IL-6) were determined via ELISA using commercial kits (EKC38669 Biomatik, EKC38664 Biomatik, and EH2IL6 Invitrogen, respectively) following manufacturers’ protocols and previously described methods^29^. Urinary norepinephrine (NE) concentrations were determined using a competitive ELISA (CatCombi Adrenaline/Noradrenaline ELISA, RE59242 Tecan Trading AG, Switzerland) as described by Ralph et al.^30^, following the manufacturer’s instructions. All samples were assayed in triplicate using a microplate reader. Final concentrations were calculated using assay-specific standard curves and, for renal tissues, adjusted for homogenate protein levels.

### Determination of kidney NE concentration

Kidneys were homogenized in 1 mL of 0.15 M perchloric acid, 0.1 mM EDTA, and 180 nM 3,4-dihydroxybenzylamine (DHBA; internal standard). Homogenates were centrifuged at 4°C for 20 min at 12,000×g. Protein content was determined in the remaining pellet by the Bradford method^31^. In the supernatant, monoamines were purified using alumina adsorption, as previously described^32^. A volume of 900 μl of supernatant was incubated for 30min in a rotatory mixer with 600 μl of 1.5 M Tris-HCl buffer (pH 8.5), 20 mM EDTA, and 10 mg alumina. After rinsing, the pellet was resuspended in 250 μl of 0.15 M perchloric acid and centrifuged at 4°C for 10 min at 12,000 g. The supernatant was filtered using a 0.22-mm filter (Millex Syringe Filter; Milipore, Burlington, MA), and 20 μl were injected into the HPLC-ED for NE measurement as previously described^33^. Separation was performed with a C-18 column (250 x 4 mm, Purospher, 5 mm; Merck, Darmstadt, Germany), preceded by a C-18 4 x 4-mm guard column. The mobile phase was composed of 100 mM sodium dihydrogen phosphate monohydrate, 10 mM sodium chloride, 0.1 mM EDTA, 0.38 mM sodium 1-octanesulfonic acid, and 6% methanol (pH 3.5), with a flow rate of 1.0 ml/min. The potential in the electrochemical detector (Decade 2; Antec Scientific, The Netherlands) was set to 0.40 V versus Ag/AgCl reference electrode.

Chromatography data were plotted with Class-VP software (Shimadzu, Kyoto, Japan), and quantification was performed by the internal standard method (DHBA) based on the peak height. All samples were measured in one assay. The intra-assay coefficient of variation of NE detection was 1.1%.

### Immunohistochemistry

Paraffin-embedded kidney sections were deparaffinized and rehydrated, followed by heat-induced antigen retrieval in Tris-EDTA buffer (pH 9.0). After blocking nonspecific binding with 2% goat serum for 30 minutes, sections were incubated overnight at 4 °C with a primary anti-CD68 antibody (1:100, v/v; ab31630, Abcam). Endogenous peroxidase activity was subsequently quenched by incubation with 3% hydrogen peroxide. Sections were then incubated with a secondary anti-mouse antibody, and immunoreactivity was visualized using 3,3′-diaminobenzidine (DAB). Nuclei were counterstained with hematoxylin. Images were acquired using a Leica light microscope and Quantimet software (Leica Biosystems). CD68-positive cells were quantified in 8–10 randomly selected high-power fields (400×) per section by an investigator blinded to group allocation using ImageJ.

### Immunofluorescence

Slides containing kidney sections were deparaffinized, rehydrated, and subjected to antigen retrieval in 10 mM citric acid buffer (pH 6.0). Sections were washed three times for 5 min in Tris-buffered saline with 0.1% Tween-20 (TBST) and incubated with 10% H₂O₂ under white light for 30 min to reduce paraffin-related autofluorescence. After additional washes in TBST, nonspecific binding was blocked with 5% goat serum for 30 min. Sections were then incubated overnight at 4°C with primary antibodies against CD68 (1:100, ab31630, Abcam) and CD206 (1:100, ab300621, Abcam). The following day, slides were washed three times in TBST and incubated for 1 h with secondary antibodies (Alexa Fluor 647 and Alexa Fluor 555; 1:500) and DAPI for nuclear counterstaining. After final washes, slides were mounted, and images were acquired at 400× magnification using the EVOS M7000 Imaging System. M2 macrophages were defined as CD68⁺/CD206⁺ double-positive cells and quantified in HF rats with or without empagliflozin treatment.

### Cell culture

The THP-1 human acute monocytic leukemia monocyte cell line (ATCC TIB-202, Manassas, VA) was obtained from the American Type Culture Collection. 200,000 cells/ml were cultured in RPMI 1640 media (GIBCO Invitrogen, Grand Island, NY) supplemented with 10% heat-inactivated fetal bovine serum (FBS, GIBCO Invitrogen, Grand Island, NY), 1% penicillin/streptomycin (GIBCO Invitrogen, Grand Island, NY), 1% sodium pyruvate (GIBCO Invitrogen, Grand Island, NY), and 0.1% 2-mercaptoethanol (Sigma-Aldrich, St. Louis, MO) at 37°C, 5% CO_2_ in a humidified tissue culture incubator. The media was replaced every 3 days, and 1×10^6^ cells/mL at passages 6-18 were seeded onto 6-well tissue culture plates for the polarization experiments. The cells were first activated with 10 ng/ml Phorbol 12-Myristate 13-Acetate (PMA, P8139 Sigma-Aldrich, St. Louis, MO) for 24h, followed by treatment with either 20 ng/ml Interferon-γ (IFN-γ, 300-04, PeproTech, Cranbury, NJ Aldrich, St. Louis, MO) and 250 ng/ml lipopolysaccharide (LPS, Sigma-Aldrich) for the M1 phenotype or 30 ng/ml Interleukin-4 (IL-4, 200-04, PeproTech) for the M2 phenotype, in the presence of vehicle (0.1% DMSO), 1 µM, or 10 µM of EMPA (catalog A12440, Adooq Bioscience, Irvine, CA) for 24h. The human kidney cell line (HK-2) was purchased from ATCC (CRL-2190) and cultured in high-glucose Dulbecco’s Modified Eagle Medium (DMEM) supplemented with 10% heat-inactivated FBS (GIBCO Invitrogen), 1% penicillin/streptomycin (GIBCO Invitrogen), 50 ng/ml bovine pituitary extract (BPE) (13028-014, Thermo Fisher Scientific, Waltham, MA), and 5 ng/ml human recombinant epidermal growth factor (hEGF) (10450-013, Thermo Fisher Scientific) at 37°C, 5% CO_2_ in a humidified tissue culture incubator. Cell media was replaced every 3 days. Cells from passages 4–12 were seeded onto 6-well tissue culture plates and grown to confluence for experiments. Cells were treated with vehicle, 1 µM EMPA, 1 µM NE (HY-13715, MCE MedChem Express, Monmouth Junction, NJ) alone or in combination with 1 µM EMPA for 48h. After the treatments, cells were washed with ice-cold phosphate-buffered saline (PBS, GIBCO Invitrogen, Grand Island, NY), lysed in 500 µl of TRIzol Reagent (Invitrogen, Waltham, MA), and stored at -80°C.

### Real-Time Quantitative PCR (RT-qPCR)

Total RNA was isolated, quantified, treated with DNase I, and reverse-transcribed into first-strand cDNA using *Super*Script III Reverse Transcription (Thermo Fisher Scientific) according to the manufacturer’s instructions. RT-qPCR was performed using SYBR Green PCR Master Mix (Thermo Fisher Scientific) on an ABI Prism 7500 Fast Sequence Detection System (Applied Biosystems, Foster City, CA). Gene expression was analyzed using the comparative threshold cycle (2^−ΔΔ^CT) method. All samples were run in triplicate. BestKeeper software was used to identify the most stable reference gene (Cyclophilin) under experimental conditions. *Tnf*, *Nos2*, *Ccr2, and Il-6* were used as markers of the M1 phenotype, whereas *Arg1*, *Cd163*, and *Mrc1* were used as markers of the M2 phenotype, as described in the literature^34,35^. The sequences of oligonucleotide primers used are listed in **Table 1**. For human proximal tubule cells, SGLT2 expression was quantified using a TaqMan Gene Expression assay (*Slc5a2,* Hs00894646_g1) and normalized to Cyclophilin A (PPIA, Hs99999904_m1) using the 2^−ΔΔCt^ method.

**Table 1.**
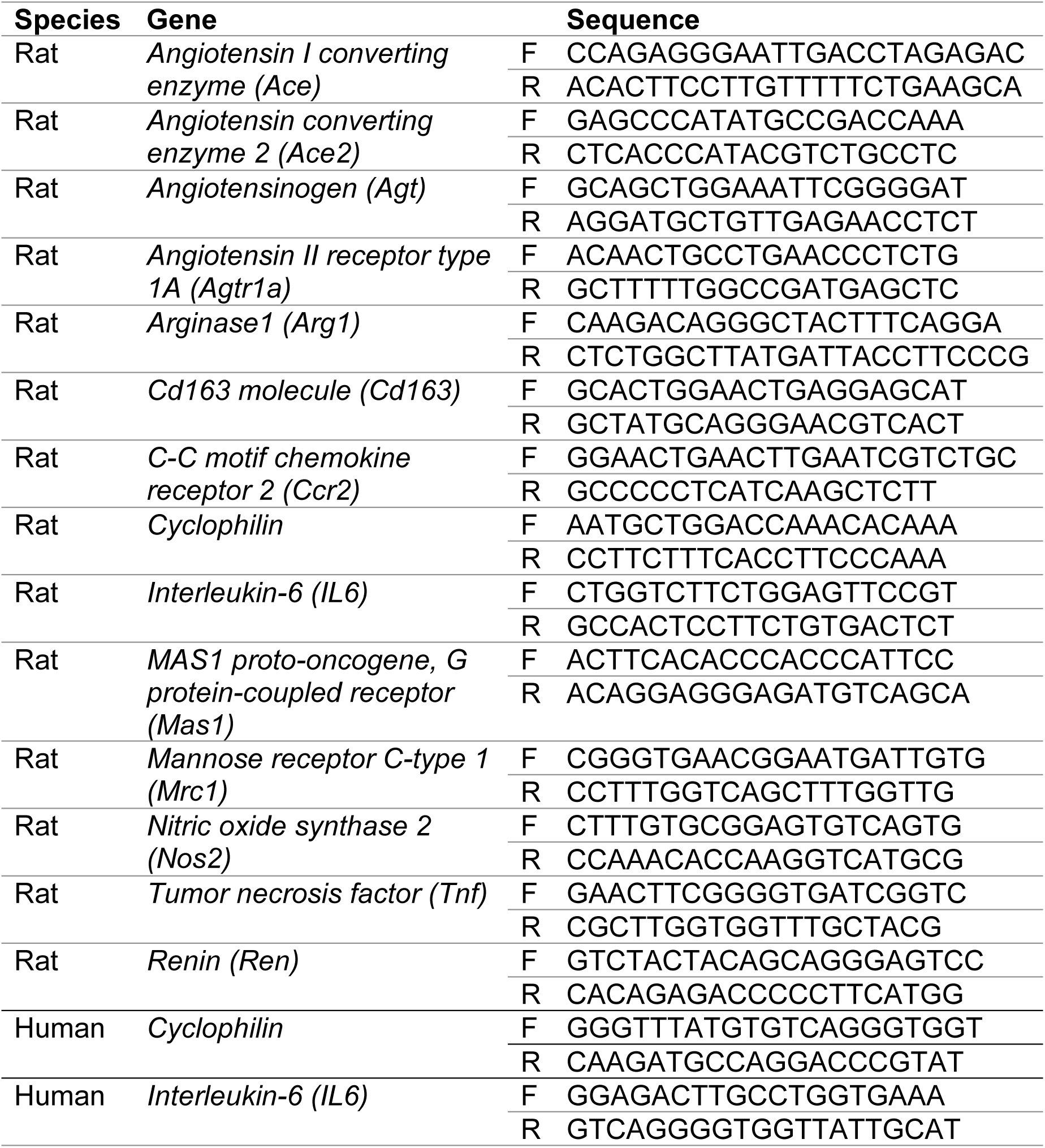
List of oligonucleotide primers.

### Statistical analysis

Data are presented as mean ± SEM. Comparisons between two groups were performed using an unpaired Student’s *t*-test. Experiments with a single independent variable were analyzed using a one-way ANOVA followed by Tukey’s post hoc test. In contrast, experiments with two independent variables were analyzed using a two-way ANOVA followed by Tukey’s post hoc test. A *P* value <0.05 was considered statistically significant.

## RESULTS

### EMPA reduces congestion and preserves GFR in HF rats despite a conserved renin-angiotensin system (RAS)

At randomization, HF was confirmed by elevated plasma BNP (Figure 1B) and reduced FAC (HF: 33±1% vs. Sham: 64±1%, P<0.001; n=32 per group). By the end of the study, EMPA reduced BNP and normalized lung water content to levels comparable to sham **(Figure 1B-C)**. Consistent with the established renoprotective effects of SGLT2 inhibitors in nondiabetic models^24,36,37^, EMPA attenuated the decline in GFR **(Figure 1D)** and limited renal mass loss **(Figure 1E)**.

To elucidate the molecular mechanisms underlying this renoprotection, we evaluated the intrarenal RAS, which is typically upregulated in HF^38^. While HF increased renal renin expression and Ang II levels, these were unaffected by EMPA treatment **(Figure 2A, C)**. Conversely, angiotensinogen was downregulated in HF and remained low regardless of treatment (**Figure 2D**). Other key components, including *Ace*, *Ace2*, *Agtr1*, and Ang (1-7) levels, remained constant across all groups (**Figure 2B, E–G**). While HF increased *MasR* expression, EMPA exerted no regulatory effect (**Figure 2H**). Collectively, these findings demonstrate that, while EMPA successfully restores renal function and alleviates congestion in HF, its protective effects are independent of modulation of the intrarenal RAS.

**Figure 2.**
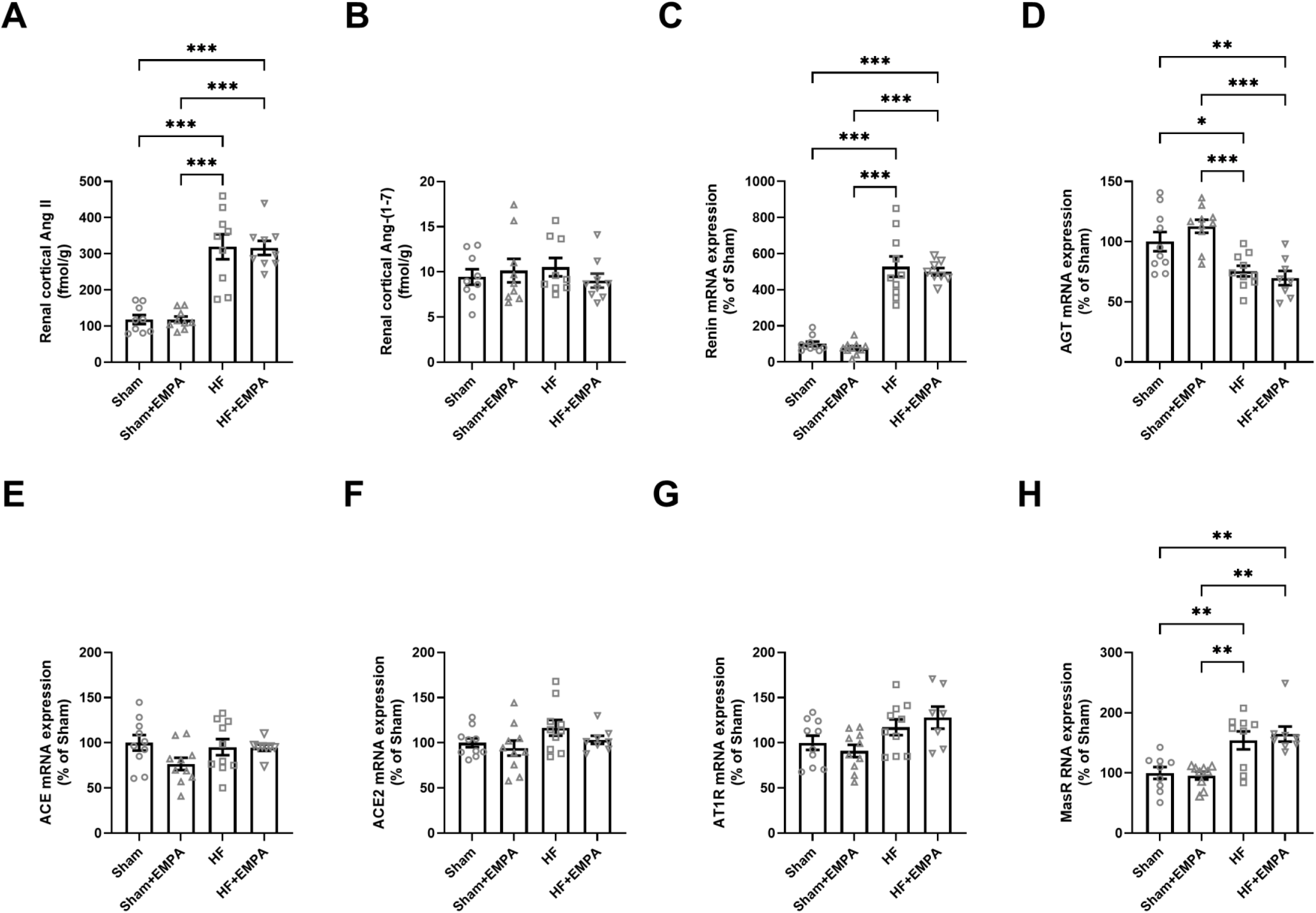
Empagliflozin does not modify the intrarenal renin–angiotensin system in heart failure. Determination of kidney **(A)** angiotensin II (Ang II) and **(B)** angiotensin-(1-7) [Ang-(1-7)] concentration. Expression analysis of renal **(C)** *renin* **(D)** Angiotensinogen (*Agt*), **(E)** Angiotensin-converting enzyme (*Ace*), **(F)** angiotensin-converting enzyme 2 (*Ace2*), **(G)** Angiotensin II receptor type 1 (*Agtr1*), and **(H)** Mas receptor (*Mas1*). Data presented as mean ± SEM (n = 7-10 rats per group). Statistical analysis was performed using two-way ANOVA followed by Tukey’s post hoc test. *P < 0.05, **P < 0.01 and ***P < 0.001.

### EMPA reduces urinary and cortical NE in HF rats, consistent with attenuated intrarenal sympathetic drive

To test the hypothesis that EMPA’s protective effects are associated with attenuation of renal sympathetic activity, we measured urinary NE excretion, a commonly used surrogate of renal sympathetic tone^39^. HF rats exhibited markedly higher NE excretion than sham rats at both randomization and the end of the treatment period, consistent with enhanced renal sympathetic activation. Empagliflozin treatment reduced urinary NE excretion by approximately 50% **(Figure 3A)**. This reduction was paralleled by intrarenal changes, with renal cortical NE content elevated in HF and reduced by ∼50% following EMPA treatment **(Figure 3B)**.

**Figure 3.**
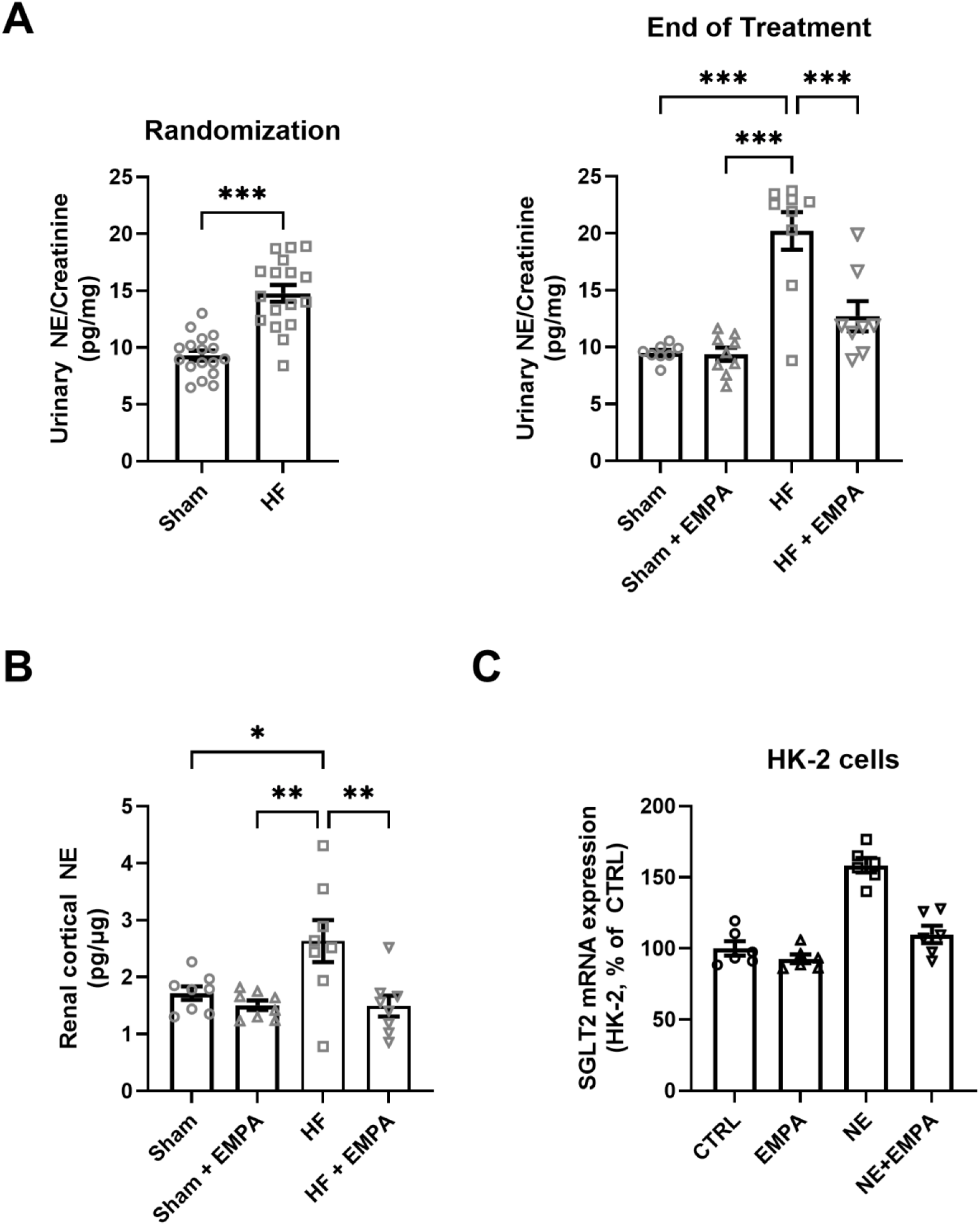
Empagliflozin reduces renal norepinephrine in heart failure and prevents norepinephrine-induced SGLT2 upregulation in proximal tubule cells. **(A)** Urinary norepinephrine (NE) excretion at randomization and posttreatment (n = 8-9 rats per group). **(B)** Renal cortical norepinephrine (NE) concentration. **(C)** Sodium-glucose cotransporter 2 (SGLT2; *Slc5a2*) mRNA expression in HK-2 human proximal tubule cells treated with vehicle, norepinephrine (NE), empagliflozin (EMPA), or NE + EMPA (n = 6). Data are presented as mean ± SEM. For *in vivo* experiments, statistical analysis was performed using an unpaired Student’s *t* test for randomization data (panel A, n = 17 rats per group) and two-way ANOVA followed by Tukey’s post hoc test (panels A and B). For *in vitro* experiments, one-way ANOVA followed by Tukey’s post hoc test was used. *P < 0.05, **P < 0.01, and ***P < 0.001.

To investigate whether EMPA directly interferes with the sympathetic regulation of SGLT2, we utilized an *in vitro* model using the HK-2 proximal tubule (PT) cell line. Building on previous findings that renal denervation prevents HF-induced SGLT2 upregulation^26^, we tested whether EMPA could directly antagonize NE-mediated induction of SGLT2. Accordingly, NE treatment significantly upregulated SGLT2 expression in HK-2 cells, and this effect was completely abrogated by the co-treatment with EMPA (**Figure 3C**). Taken together, these findings indicate that EMPA reduces urinary and renal cortical norepinephrine levels in HF, consistent with attenuation of intrarenal sympathetic activity. In parallel, EMPA prevents NE-induced SGLT2 upregulation in proximal tubule cells, supporting the existence of a renal sympathetic-SGLT2 interaction.

### EMPA promotes a phenotypic shift from pro-inflammatory M1 to anti-inflammatory M2 macrophages in HF kidneys

To investigate the downstream consequences of EMPA-mediated NE reduction, we evaluated the renal inflammatory response. Histological staining for the pan-macrophage marker CD68 revealed that total macrophage abundance was higher in HF rats compared with sham controls, with no difference between EMPA-treated and untreated HF groups. (**Figure 4**). However, gene expression profiling suggested that EMPA significantly affected macrophage polarization. In HF rats, we observed a predominantly pro-inflammatory M1-like profile, characterized by the upregulation of *Tnf*, *Nos2*, *Ccr2*, and *Il-6,* while the M2-like markers *Arg1*, *Mrc1*, *and CD163* remained at sham levels. EMPA treatment successfully reversed this signature: expression of M1 markers was significantly attenuated, while M2 markers were markedly upregulated (**Figure 5A-G**). This phenotypic shift was confirmed via dual-immunofluorescence staining for CD68 and the M2-specific marker CD206 (encoded by *Mrc1*). In HF kidneys, CD68^+^ macrophages largely lacked CD206 expression; conversely, in HF EMPA kidneys, the majority of the macrophage population was double labeled for CD206 (0.7 ± 0.3 vs. 4.7 ± 1.2 CD68⁺/CD206⁺ double-positive cells per field, P < 0.03, n = 3 rats/group) (**Figure 5H-I**). These results demonstrate that while EMPA does not alter total macrophage recruitment to the kidney in HF, it shifts the local immune environment from a pro-inflammatory M1 state toward a reparative M2-like phenotype.

**Figure 4.**
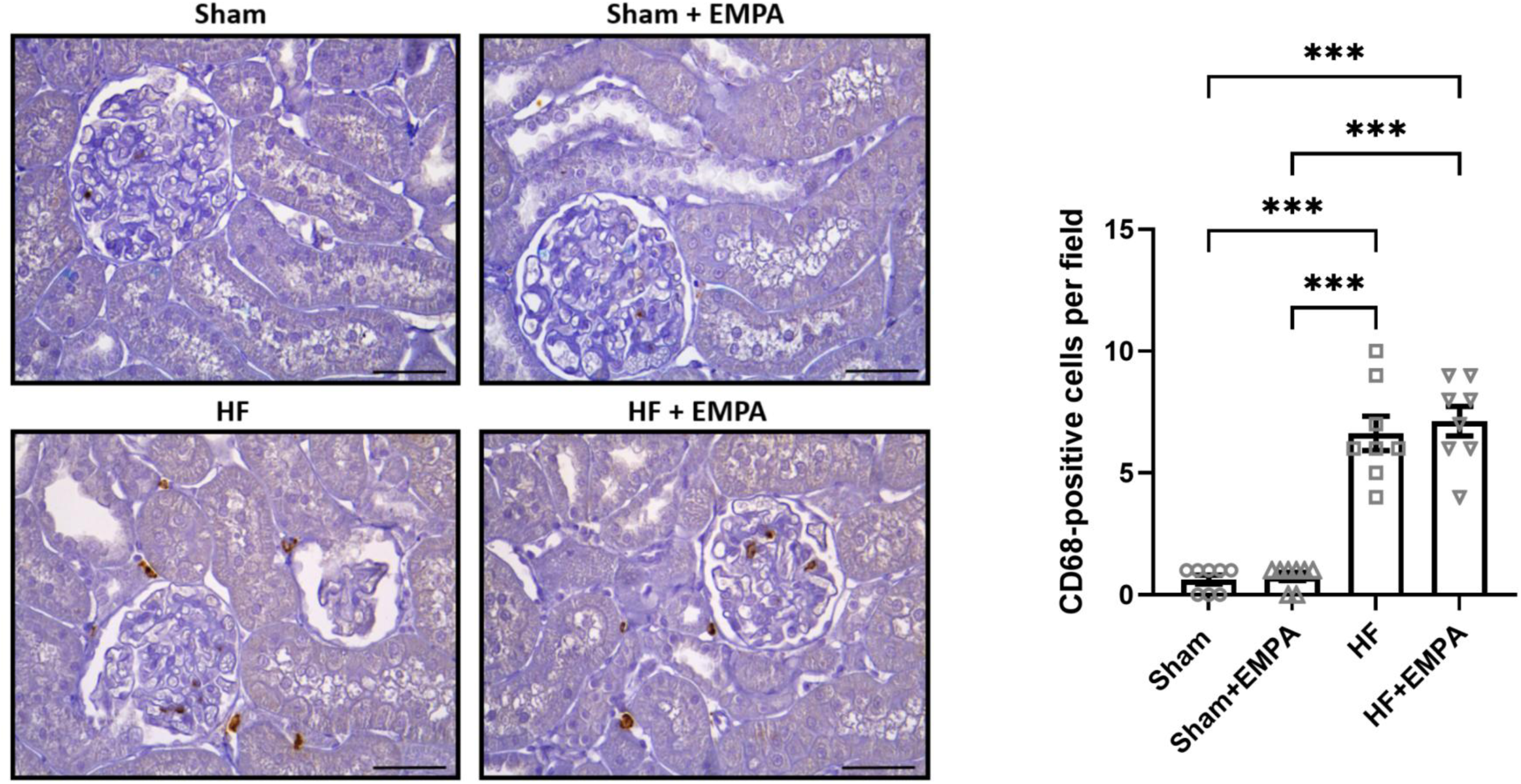
Empagliflozin does not alter total renal macrophage abundance in heart failure. Representative immunohistochemical staining for the pan-macrophage marker cluster of differentiation 68 (CD68) in renal cortical sections. Scale bar = 50 µm. Quantification of CD68^+^ cells. Data are presented as mean ± SEM (n = 9 rats per group). Quantification was performed in 8–10 random high-power fields per section. Statistical analysis was performed using two-way ANOVA followed by Tukey’s post hoc test. ***P < 0.001.

**Figure 5.**
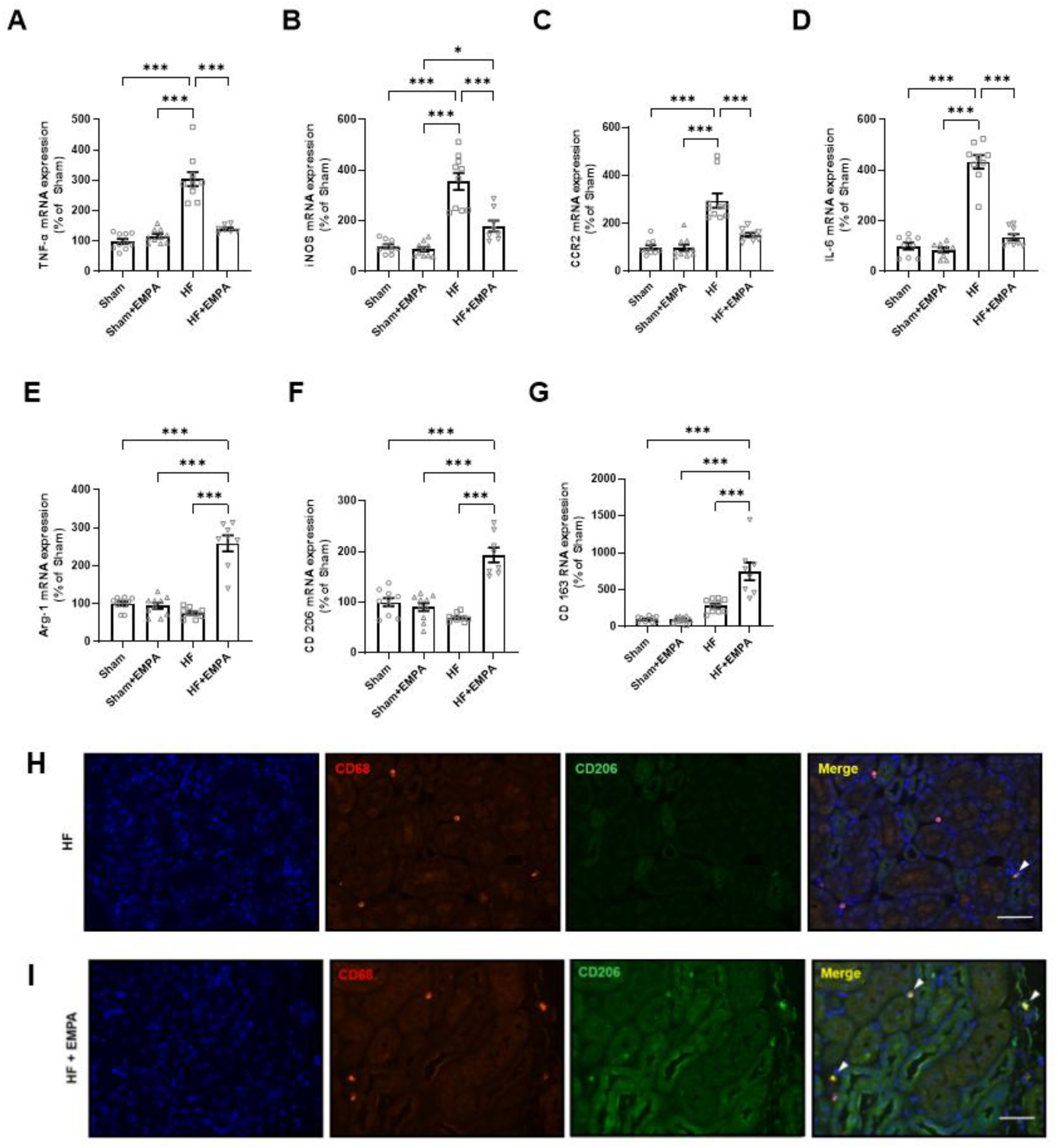
Empagliflozin favors a shift from pro-inflammatory M1 to anti-inflammatory M2 macrophage markers in the kidneys of rats with heart failure. Renal cortical mRNA expression of M1 macrophage markers (A) *Tnf*, (B) *Nos2*, (C) *Ccr2*, and (D) *Il-6*, and M2 macrophage markers (E) *Arg1*, (F) *Mrc1* (CD206), and (G) *Cd163* in sham rats and heart failure (HF) rats treated or not with empagliflozin (EMPA). Data presented as mean ± SEM (n = 7-10 rats per group). Statistical analysis was performed using two-way ANOVA followed by Tukey’s post hoc test. *P < 0.05 and ***P < 0.001. **(H-I)** Representative dual-immunofluorescence images of renal cortical sections from HF rats treated or not with empagliflozin (EMPA), stained for CD68 (pan-macrophage marker, red) and CD206 (M2 marker, green), with nuclei counterstained with DAPI (blue). CD68⁺/CD206⁺ double-positive macrophages are identified by yellow signals in merged images and are indicated by white arrows. Scale bar = 50 µm.

### EMPA does not directly influence macrophage polarization *in vitro*

To determine whether the observed shift toward an M2-like phenotype reflected a direct action of EMPA on macrophages, we utilized the THP-1 monocyte cell line. We first assessed whether EMPA alone could induce polarization in the presence of the activator PMA. Under these baseline conditions, EMPA treatment (1 mM or 10 mM) failed to alter the expression of either *Tnf* or *Mrc1*, indicating that EMPA lacks intrinsic polarizing activity (**Figure 6A**). We next investigated whether EMPA could interfere with established polarizing stimuli. To induce an M1-like phenotype, THP-1 cells were treated with Interferon-γ (IFN-γ) and Lipopolysaccharide (LPS). EMPA treatment had no impact on this process, as *Tnf* expression was upregulated to the same extent across all groups (**Figure 6B**). Similarly, EMPA failed to affect M2-like polarization induced by Interleukin-4 (IL-4), with *Mrc1* expression levels remaining unchanged across drug concentrations (**Figure 6C**). These findings demonstrate that EMPA does not directly influence macrophage polarization. Consequently, the M2-like shift observed in HF EMPA rats likely results from the drug’s effects on the renal microenvironment (e.g., reduced sympathetic activity), rather than from a direct modulation of macrophage polarization.

**Figure 6.**
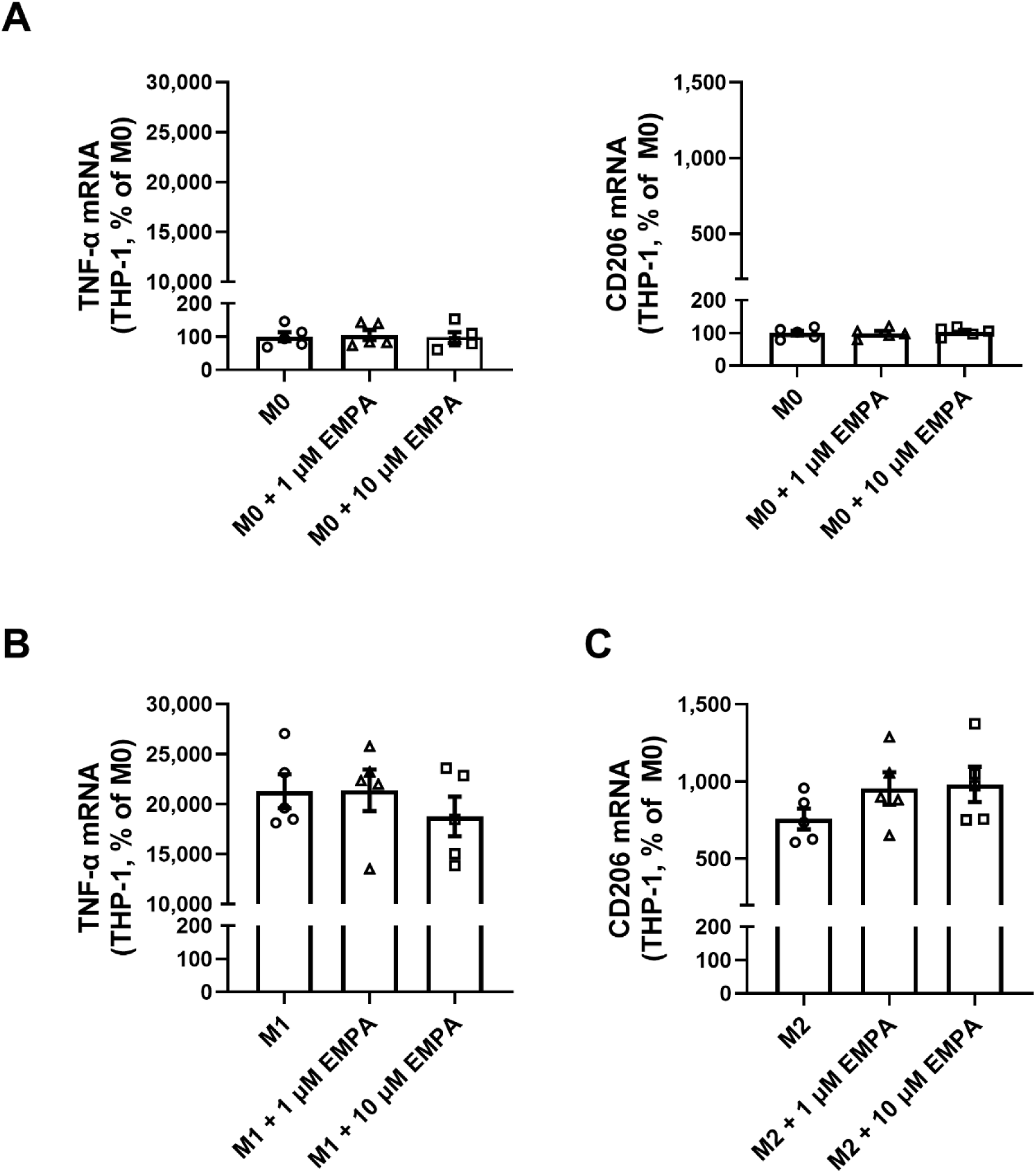
Empagliflozin does not directly alter macrophage polarization markers in THP-1 cells. **(A)** THP-1-derived macrophages under basal (M0) conditions were treated with empagliflozin (EMPA; 1 or 10 μM), and mRNA expression of TNF-α and CD206 was assessed. **(B)** THP-1 macrophages polarized toward a pro-inflammatory phenotype (M1) and treated with EMPA (1 or 10 μM); TNF-α mRNA expression is shown. **(C)** THP-1 macrophages polarized toward an alternative phenotype (M2) and treated with EMPA (1 or 10 μM); CD206 mRNA expression is shown. Gene expression was normalized to cyclophilin and expressed relative to M0 controls. Data are presented as mean ± SEM; individual data points represent five independent experiments performed in duplicate.

### EMPA antagonizes NE-induced IL-6 and TNF-α expression in PT cells

Considering our findings that EMPA does not directly influence macrophages, we investigated whether it modulates the renal microenvironment that drives polarization. Interleukin-6 (IL-6) is a known driver of M1 polarization, reinforcing the expression of pro-inflammatory cytokines like TNF-α. Since NE induces IL-6 in PT cells^40^, we hypothesized that EMPA might indirectly promote M2 polarization by blocking NE-mediated cytokine production in PT cells. Thus, we measured IL-6 expression and secretion in HK-2 cells. While EMPA alone did not affect baseline IL-6 levels, NE increased both IL-6 mRNA expression and protein secretion by approximately 50% (**Figure 7A-B**). This upregulation was completely mitigated by EMPA co-treatment, maintaining IL-6 at control levels. Collectively, these results suggest that EMPA’s renoprotective effects and its ability to shift macrophage polarization are, at least in part, mediated by antagonizing the NE-induced pro-inflammatory secretome of the proximal tubule.

**Figure 7.**
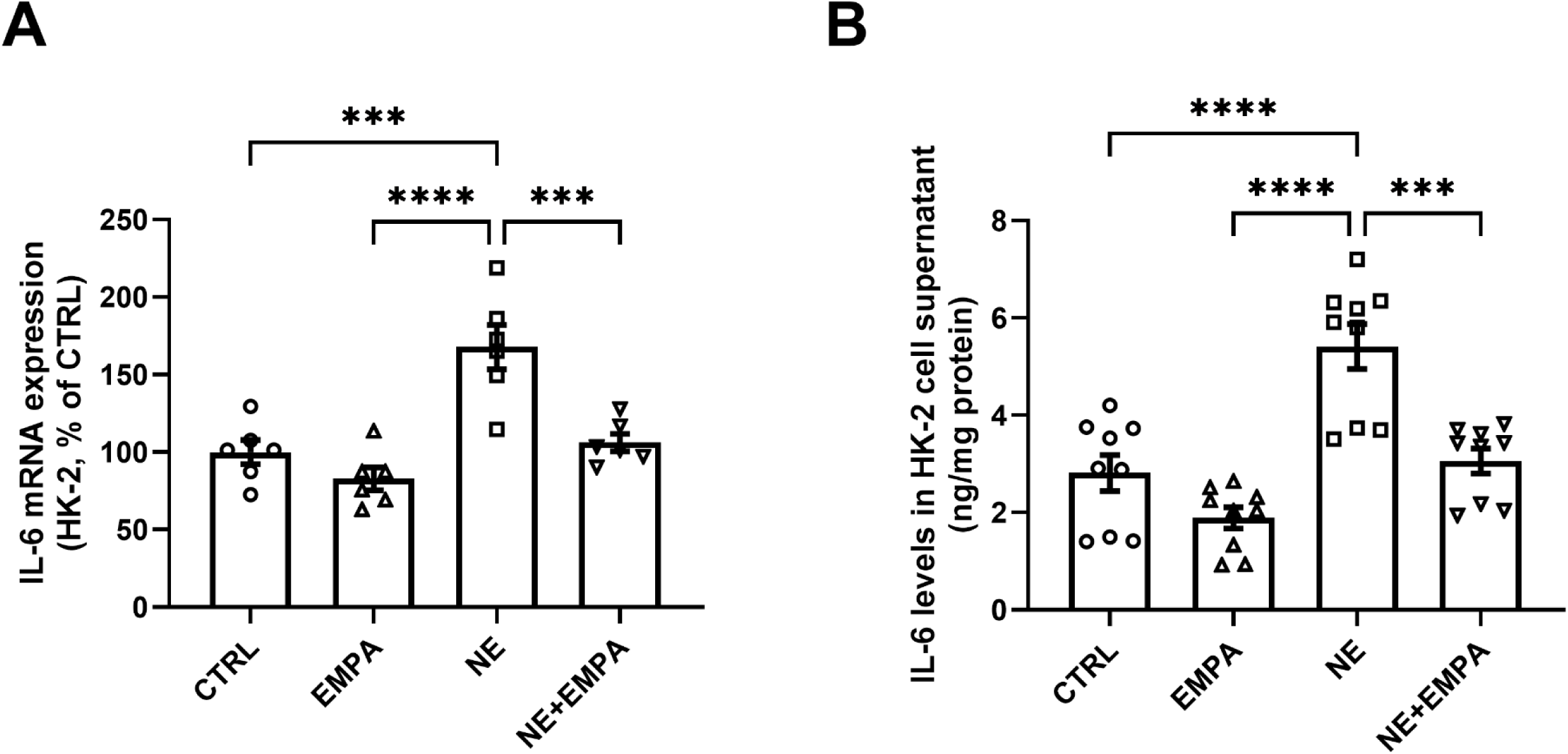
Empagliflozin antagonizes norepinephrine-induced inflammatory responses in human proximal tubule cells (HK-2 cells). **(A)** IL-6 mRNA expression in HK-2 cells treated with vehicle (Control), empagliflozin (EMPA, 1 μM), norepinephrine (NE, 1 μM), or combined NE + EMPA. IL-6 mRNA levels were normalized to cyclophilin and expressed as a percentage of control. Each data point represents five independent experiments performed in duplicate. **(B)** IL-6 concentration in the supernatant of HK-2 cells under the same experimental conditions, measured by ELISA and normalized to protein content. Individual data points represent nine independent experiments performed in duplicate. Data are presented as mean ± SEM. Statistical analysis was performed using one-way ANOVA followed by Tukey’s post hoc test. ***P < 0.001 and ****P < 0.0001.

## DISCUSSION

While the sympatholytic effects of SGLT2 inhibitors are increasingly recognized, the intrarenal neurohumoral pathways through which these agents act in HF remain elusive. This study provides novel evidence that EMPA targets the renal sympathetic axis, normalizing renal cortical NE content and urinary excretion independently of the intrarenal RAS. Notably, we identify a renal neuro-epithelial-immune axis associated with renoprotection. By attenuating NE-driven SGLT2 upregulation and proximal tubular inflammatory signaling, EMPA creates a microenvironment that favors a shift from a pro-inflammatory (M1) to a reparative (M2) macrophage phenotype. Collectively, our data suggest that the clinical success of SGLT2is in multidrug HF regimens (often already including RAAS blockade) may reflect their ability to modulate a neuro-inflammatory niche that classical neurohumoral agents do not directly target. Given that sympathetic activation and renal congestion are common to both reduced and preserved ejection fraction phenotypes, targeting this renal neuroinflammatory axis may help explain the consistent benefits of SGLT2i across the HF spectrum.

In HF, the sympathetic nervous system and RAAS are viewed as mutually reinforcing systems: sympathetic activation typically stimulates the juxtaglomerular apparatus via adrenergic receptors to release renin, which then cascades into Ang II production^41^. However, we found that EMPA-mediated sympatholysis occurred without altering renal renin or Ang II levels. These findings align with recent evidence from salt-sensitive^42^ and neurogenic ^43^ hypertension models in which dapagliflozin improved the pressure-natriuresis response and reduced blood pressure independently of the systemic or renal RAAS. Together, these data support the concept that SGLT2 inhibition can attenuate renal sympathetic activity through mechanisms distinct from classical RAAS suppression.

A potential mechanistic basis for this dissociation lies in the crosstalk between adrenergic signaling and SGLT2 expression in the renal proximal tubule. Our *in vitro* studies demonstrate that NE directly upregulates SGLT2 in HK-2 cells, consistent with previous reports ^44,45^. Increased SGLT2 activity enhances sodium-glucose reabsorption, promoting tubular stress and IL-6 production. In the context of HF, where sympathetic tone, SGLT2 expression, and inflammatory signaling are simultaneously elevated, this interaction may establish a maladaptive positive feed-forward loop in which NE amplifies epithelial inflammatory signaling, thereby sustaining a pro-inflammatory microenvironment that could reinforce renal afferent excitation. Importantly, we do not propose that SGLT2 directly regulates NE release under physiological conditions. Rather, we suggest that in the heart failure kidney, SGLT2-dependent epithelial stress may contribute to the maintenance of sympathetic overactivity through reno-neural mechanisms. This context dependence likely explains why EMPA reduced renal NE levels in HF but not in sham animals, in which this neuro-epithelial-inflammatory axis is not engaged. By attenuating sympathetic-driven epithelial stress, EMPA may help prevent the transition from functional impairment to structural remodeling, a hallmark of CRS. This interpretation is consistent with the preservation of GFR observed in our model and with reports of reduced IL-6 levels and enhanced catecholamine metabolism in other SGLT2i-treated inflammatory states⁴⁶.

In line with findings in the 5/6 nephrotomy model of chronic kidney disease (CKD)^47^, EMPA did not alter total macrophage abundance but shifted polarization M1 toward the M2 macrophage phenotype. Similar immunometabolic reprogramming has been observed in experimental models of myocardial infarction and obesity^48,49^. In diet-induced obesity, SGLT2is reduced M1 macrophage accumulation while promoting M2 polarization in adipose tissue and the liver, lowering TNF-α and improving insulin sensitivity^49^. Our findings extend this paradigm to nondiabetic HF, where the kidney, analogous to adipose tissue in metabolic disease, becomes a site of chronic neuro-inflammatory stress. By dampening sympathetic-driven tubular stress, EMPA may create conditions that favor reparative macrophage signaling and inflammation resolution.

Our in vitro findings support an indirect immunomodulatory mechanism. While some studies using supra-therapeutic doses (80 μM)^50^ or murine macrophages^51,52^ suggest a direct anti-inflammatory action of EMPA, our results at clinically relevant concentrations (1-10 μM) suggest that macrophage reprogramming in vivo reflects modification of the renal microenvironment rather than intrinsic macrophage effects. Alternatively, the dissonance between our data and the literature might result from the differential systemic environments (Influenza virus infection^50^ or high glucose^52^) that may not reflect the nondiabetic HF environment.

This study has limitations. A direct assessment of renal sympathetic nerve activity, either via renal nerve recordings or renal NE spillover measurements, was not performed. Thus, sympathetic tone was inferred from surrogate indices, including urinary excretion and renal cortical NE content, which may be influenced by processes related to neurotransmitter release, reuptake, metabolism, and excretion. In addition, the conclusion that EMPA promotes macrophage repolarization indirectly by affecting the renal microenvironment is supported by the absence of direct effects on THP-1 macrophage polarization and by concomitant reductions in renal NE levels and PT inflammatory signaling. Although causality between sympathetic attenuation, reduced tubular cytokine production, and macrophage repolarization cannot be definitively established from the present data, the concordant in vivo and in vitro findings consistently support attenuation of renal sympathetic-inflammatory signaling as a central mechanism of EMPA action in HF.

In summary, EMPA-mediated attenuation of renal sympathetic activity, occurring independently of detectable intrarenal RAAS modulation, contributes to renoprotection by disrupting a maladaptive neuro-epithelial-immune axis. Beyond these intrarenal effects, modulation of renal sympathetic tone may also influence kidney-heart communication. Given the established role of renal afferent signaling in sustaining central sympathetic outflow, attenuation of renal neuroinflammatory stress may indirectly dampen systemic sympathoexcitation and contribute to cardioprotection. Although neural signaling was not directly interrogated, our findings support a model in which SGLT2 inhibition confers additive clinical benefit by targeting a renal neuroinflammatory niche that amplifies cardiorenal dysfunction.

## DATA AVAILABILITY

Data supporting the findings of this study are available from the corresponding author upon reasonable request.

## NON-STANDARD ABBREVIATIONS AND ACRONYMS

EMPA: Empagliflozin
HF: Heart Failure
NE: Norepinephrine
RAS: Renin-Angiotensin System
SGLT2is: SGLT2 Inhibitors
SNS: Sympathetic Nervous System

